# The human sperm beats anisotropically and asymmetrically in 3D

**DOI:** 10.1101/795245

**Authors:** Hermes Gadêlha, Paul Hernández-Herrera, Fernando Montoya, Alberto Darszon, Gabriel Corkidi

## Abstract

The canonical beating of the human sperm flagellum is postulated to be symmetric. This is despite the reported asymmetries inherent to the flagellar axonemal structure, from distribution and activation of molecular motors to, even, the localisation of regulatory ion channels. This raises a fundamental question: how symmetric beating is possible within such intrinsically asymmetric flagellar complex? Here, we employ high-speed 3D imaging with mathematical analysis capable of resolving the flagellar movement in 4D (3D+time). This reveals that the human sperm beating is both anisotropic and asymmetric, and composed by a superposition of two transversal waves: an asymmetric travelling wave and a symmetric standing wave. This novel anisotropic travelling-pulsation mechanism induces sperm rolling self-organisation and causes a flagellar kinematic illusion, so that the beat appears to be symmetric if observed with 2D microscopy. The 3D beating anisotropy thus regularises the intrinsic flagellar asymmetry to achieve symmetric side-to-side movement and straight-line swimming.

---

Recent precision cryo-electron tomography of individual dyneins in sea-urchin sperm flagellum unveiled an asymmetrical distribution of molecular motors [1, 2]. Asymmetric waveforms have also been reported in mammalian sperm, such as murine spermatozoa [3, 4], while an asymmetric beating mode is a typical feature in the *Chlamydomonas* flagellum [5–9]. Whilst structural anisotropies are inherent to the mammalian sperm architecture [10–12], other sources of asymmetry are equally present at the molecular level, from asymmetric dynein activity [1, 2, 13–15], to an ever growing number of asymmetric regulatory complexes [6, 8, 16–20], including anisotropically localised membrane ion channel along the flagellum [17]. This is further augmented by external sources of asymmetry, such as the hydrodynamic influence of solid surfaces during for sperm boundary accumulation near to the coverslip [21–23]. This raises a fundamental question: given this multilayered hierarchy of distinct asymmetries that are intrinsically present, how does the human sperm flagellum achieve symmetric beating and straight-line swimming trajectories typically observed under the microscope [10, 24–28]?

Indeed, our understanding of the human sperm flagellum beating is centred on the dogma of symmetric beating [29–31]. This have been postulated based on the wealth of observations thus far showing straight-line swimming trajectories for sperm beating symmetrically, as well as symmetric travelling waves of curvature of the waveform [10, 24, 26–28, 32]. As such, beating symmetry is the cornerstone of any comparative study on flagellar undulation [24, 29, 32], especially so while assessing symmetry-breaking events, such as in sperm capacitation, hyperactivation and buckling phenomena [18, 19, 32–34]. Theoretical and computational models equally, and ubiquitously, invoke symmetry, ranging from flagellar kinematic models to structural mechanics and even motor control hypothesis [5, 10, 12, 26, 35–39]. How to even reconcile symmetric swimming observations with the umbrella of asymmetries that are present at the molecular level in the human spermatozoa, is still open to debate.

This scenario is aggravated by the fact that the literature to date has been predominately limited to planar projections of the flagella beat [10, 24–31], albeit the human sperm flagellum beats intrinsically in three-dimensions. Despite numerous efforts to capture the flagellar beat in 3D [34, 40–52], the importance of threedimensional component of the beat is yet to be fully recognised at the human fertility realm [53]. Furthermore, the rapid movement of the flagellar beating in 4D (3D+time) continues to challenge recent advances in video-microscopy and holography techniques [34, 40–42, 46–50], and thus still remains as a major bottleneck for any automated mathematical image processing algorithm [*ibid*].

Here we employ a high-precision rapid capture of the human sperm flagellum in 4D. This revealed that the flagellar beating is both *asymmetric* and *anisotropic*, and composed by a superposition of an asymmetric travelling wave and a symmetric standing wave in the transverse direction. This novel beating anisotropy regulates the sperm rolling self-organisation and induces a flagellar kinematic illusion, in which the flagellar beat appears to be symmetric if observed with 2D microscopy. This impacts the modulation of curvature in 3D, the waveform torsion and the beating chirality, which is characterised by a new kinematic flagellar perversion phenomenon. Our observations reconcile the growing number of studies reporting anisotropic and asymmetric regulatory complexes within the flagellar scaffold with the canonical observations of symmetric beating and straight-line swimming. The latter unearth a subtle anisotropic control mechanism capable of regularising intrinsic flagellar asymmetries to recover symmetry for the purpose of sperm transport.

## RESULTS

### The non-slip boundary is a weak source of asymmetry to the flagellar beat

The rapid movement of human sperm flagella was recorded with high spatiotemporal resolution in 3D, Fig. 1(a,b), as described in the Methods section. We investigate the hydrodynamic impact of a nearby coverslip on the beating symmetry, given the uneven influence of hydrodynamic forces due to the presence of solid boundary, i.e. non-slip surface [22, 23, 26]. Fig. 1(a,b) shows, respectively, the flagellar beating of a spermatozoon swimming near and far from coverslip. Spermatozoa displayed bi-directional rolling around their swimming axis, though the majority of cells (28 cells) rolled counterclockwise when viewed from the anterior end (arrows in Fig. 1(c,d)), only two cells rolled clockwise, see SI text and SI-Fig 3. The combined rolling and translation motion of the sperm flagellum leads to helical trajectories of the mid-flagellar point, red tracers in Fig. 1(a,b). The projection of the mid-flagellar trajectory, red curves in Fig. 1(c,d), known as flagelloids [51, 52], displayed a bewildering array of geometrical patterns, from rotating star-shapes to triangles, squares and looping patterns with polar symmetry, SI-Fig. 3.

**FIG. 1.**
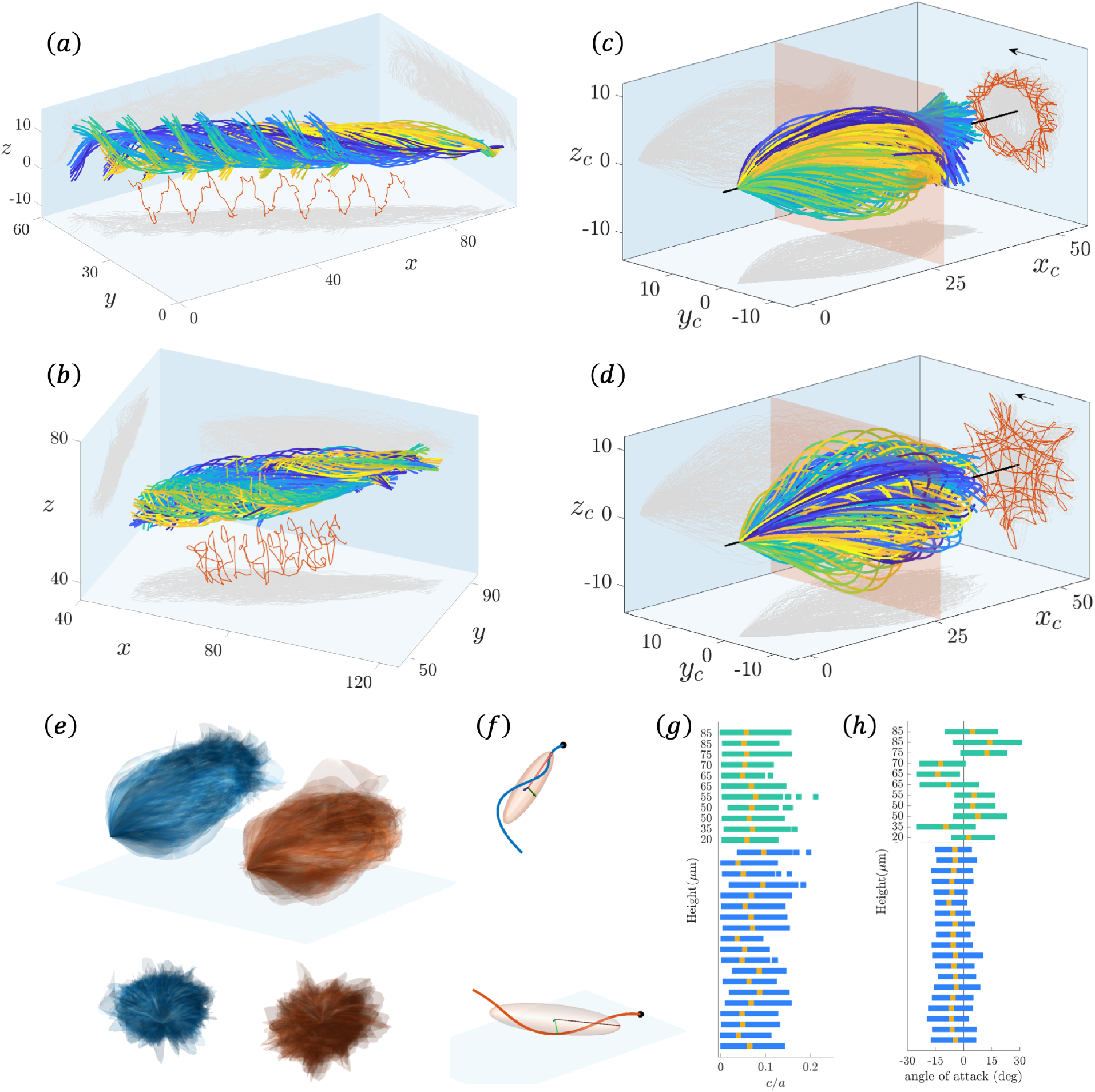
Human sperm flagellar beating near (a,c) and far from the coverslip (b,d) in 3D: (a,b) Flagellar waveform relative to the laboratory fixed frame of reference (*x,y,z*) and (c,d) relative to the comoving frame of reference (*x_c_, y_c_, z_c_*) (Methods section). In (a-d) time increases from blue to yellow for each period, red curves depict the trajectory of the mid-flagellar point, indicated by the red plane cross-section in (c,d). Light grey curves in (a-d) plot the flagellar projection in each plane. Note that 2D microscopy only captures the planar xy-projection depicted. The flagellum rotates around the mean swimming axis, i.e the rolling axis, depicted by the black straight-line in (c,d) with the arrow indicating the rolling direction. (c,d) show highly symmetric beating in both planar (*xy*) and out-of-plane direction (*z*), see also Supporting Information video of (a,b). (e) compares the 3D flagellar envelop for sperm near (blue) to and far (red) from the coverslip, showing boundary modulation of the waveform. (f-h) The flagellar Inertia ellipsoid capturing the three-dimensionality of the beat as described in the Methods section: (f) Inertia ellipsoid and its ellipsoidal axes at a given instant for the spermatozoa in (a) and (b), respectively, in red and blue. (g) The variation of ratio between the minor and major axis of the inertia ellipsoid *c/a* at each instant for each individual sperm. Sperm near to and far from coverslip are shown, respectively, by the blue and green bars, the yellow marker shows the temporal average. If *c/a* = 0 the waveform is a purely planar curve. The flagellar beat oscillates between a planar and a weakly non-planar beat. (h) The orientation of the major axis of the inertia ellipsoid relative to the coverslip at *z* = 0, as shown by the red vector in (f), defines the angle of attack of the flagellum at each instant, negative (positive) angles for sperm swimming towards to (away from) the coverslip, with colour scheme as in (g).

At the comoving frame of reference (Methods), Fig. 1(c,d) and SI-Fig. 2, the flagellar beat is highly symmetrical in both planar (*xy*) and out-of-plane (*z*) projections, depicted by the light grey curves. Indeed, the 3D wave envelop in Fig. 1(e) has a rounded bullet-shape, and not the conical envelop commonly hypothesised [19, 54, 55]. Spermatozoa swimming far from the surface have a more symmetric flagellar envelop (red), whilst for cells near to the coverslip (blue) the bullet-shape envelop is “bent” towards the surface (top row), resembling a bird’s beak (Fig. 1(e)). The nearly oval cross-section of the wave-envelop for the cells near to the coverslip deviates weakly from the approximately circular shape for the cells far from coverslip, bottom row in Fig. 1(e). The hydrodynamic modulation of the flagellum by coverslip thus breaks weakly the symmetry of the beat in 3D. Nevertheless, the observed ratio of the volume of the 3D wave-envelop between cells swimming near to and far from the coverslip was *V*_near_/*V*_far_ = 0.75.

The 3D waveform at each instant oscillates between a purely planar and weakly non-planar beating, Fig. 1(f,g), in agreement with earlier observations [34]. The average ratio between the minor and major axis of the flagellar inertia ellipsoid for the cells “near to” and “far from” the coverslip (Methods) appears to be conserved across the population with, respectively, *c/a* = 0.0623, 0.0631, characterised by flat ellipsoids. Thus only a small non-planar component of the waveform can induce large flagellar excursions in 3D, Fig. 1(c,d). The proximity to the coverslip does not appear to influence the weak non-planarity of beat, as show in Fig. 1(g). The swimming angle of attack, however, is regulated by the proximity to the coverslip [22, 23], as shown in Fig. 1(h). Spermatozoa far from the coverslip are able to swim both towards to and away from the boundary, whilst all spermatozoa swimming near to the coverslip displayed an conserved angle of attack of −7°, directed towards the surface, Fig. 1(h). Interestingly, whilst the angle of attack oscillates around its mean, the amplitude of oscillation allows sign changes over the course of time even for the sperm swimming near to the surface. This indicates that the sperm flagellum is directed away from the coverslip for short periods when swimming close to the boundary, blue bars in Fig. 1(h).

### The 3D beating of the human sperm flagellum is anisotropic and asymmetric

The true nature of the flagellar beat in absence of all swimming translations and rotations, and thus unbiased by the sperm rolling motion, is shown in Fig. 2(b), SI-Fig. 4 and SI videos. The comoving-rolling frame of reference (Methods), Fig. 2(e), reveals that the flagellar beat is anisotropic: the wave characteristics in each transversal direction, b-plane (blue plane) and z-plane (red plane), differ dramatically. Furthermore, the beating is highly asymmetric at the b-plane, and characterised by a distinctive broken left-right symmetry resembling a ‘C’ shape, Fig. 2(e). This is in contrast with the remarkably symmetric patterns observed at the co-moving frame of reference in Fig. 2(a).

**FIG. 2.**
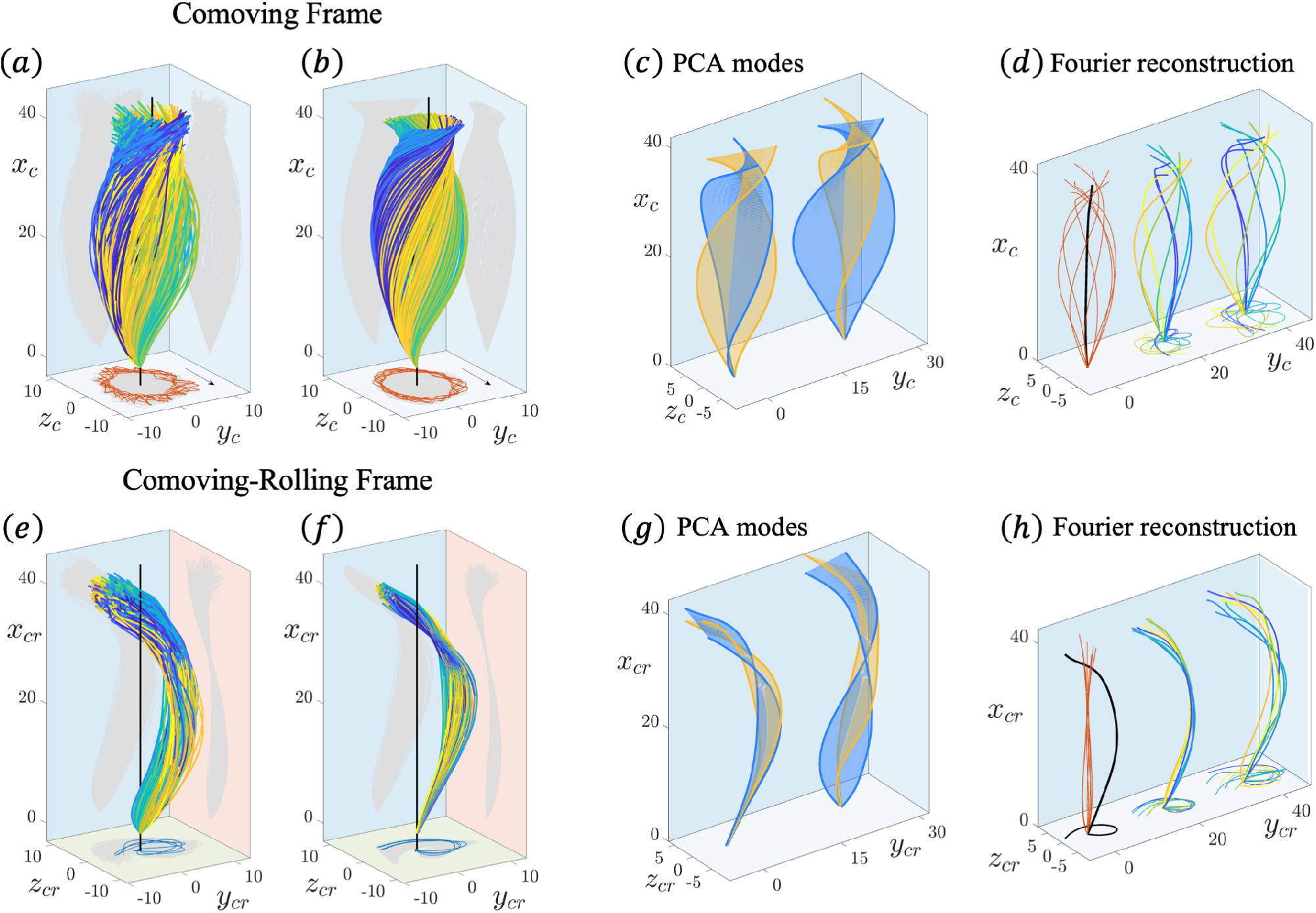
The human sperm flagellum beats anisotropically and asymmetrically in 3D. Top-row (a-d) the flagellar waveform relative to the comoving frame of reference (*x_c_, y_c_, z_c_*) for the same spermatozoon shown in Fig. 1(a,c), displaying the rolling-axis up-right (black curve) with arrow showing the rolling direction in (a,b). Bottom-row (e-h) flagellar waveform relative to the comoving and rolling frame of reference (*x_cr_, y_cr_, z_cr_*), i.e. absence of flagellar rotation around the rolling-axis, black curve in (e,f). The beating anisotropy and asymmetry is revealed at the frame of reference that moves and rotates with the flagellum. SI video shows the three changes in referential employed: fixed, comoving and comoving-rolling frames of reference. (e) introduces the three perpendicular beating planes of comoving-rolling frame of reference, denoted by “b-plane” (blue plane), which captures the planar motion of the flagellum at the *xy*-projection, the “z-plane” (red plane), responsible for the out-ofplane motion, the “rolling plane” (green plane), defined by the *yz*-plane perpendicular to the rolling axis. Second-column (b,f) show the principal component analysis (PCA) reconstruction of the waveform using the first two PCA modes. Third-column (c,g) shows the surfaces generated by the 1st PCA mode (blue) and 2nd PCA mode (orange) in the course of time, for two different views (left and right plots in (c) and (g)). The PCA reconstruction in (b,f) is a superposition of the PCA modes in (c,g). Fourth-column (d,h) display the Fourier analysis of the waveform. For both (d) and (h): left-plot shows the static (black) and dynamic (red) Fourier modes; the middle-plot shows Fourier reconstruction of the waveform using the first two modes, i.e. a superposition of the static mode (black) and dynamic mode (red); and the right-plot compares with the original data. In (a,b,d) and (e,f,h) time increases from blue to yellow for each period. The transformations between frame of references, principal component and Fourier analyses are detailed in Methods.

In both comoving and comoving-rolling frames of reference, the 3D waveform is well characterised with only two principal component modes, Fig. 2(b,f) and SI-Figs. 5,6, referred as PCA modes (Methods). In Fig. 2(c) the first two PCA modes have an identical in shape, up to a 90° rotation (compare the blue and orange surfaces), thus capturing the streamlined helical shape caused by the sperm rolling. In Fig. 2(g), however, the intrinsic asymmetric ‘C’ shape is fully detected by the 1st PCA mode alone (blue surface). The 2nd PCA mode in Fig. 2(g) (orange surface) contributes with small deviations perpendicular to the 1st PCA mode (blue surface), indicating that the waveform may be decomposed in two independent transversal directions.

Fig. 2(d,h) shows the Fourier reconstruction (Methods) using the static (black) and dynamic modes (red). The static mode captures any shape incongruence of the waveform, thus the black straight-line in Fig. 2(d) reflects the beating symmetry in both *xy* and *xz* planes. In contrast, at the comoving-rolling frame in Fig. 2(h), the static mode is characterised by a large amplitude asymmetric C-bend. The dynamic mode at the comoving frame (red curves) in Fig. 2(d) has a large amplitude and is highly symmetric in both *xy* and *xz* planes due to sperm rolling. In contrast, the dynamical mode (red curves) at the comoving-rolling frame, Fig. 2(h), has a reduced waveform amplitude and, instead, a preferential direction of motion (black curve in Fig. 2(a)). The Fourier reconstruction of the waveform is thus given by the superposition, or sum, of the static and dynamic modes, depicted by the middle-plots in Fig. 2(d,h), and agrees well with the original observations (right-plots in Fig. 2(d,h)).

### The flagellar beat is a transversal superposition of a travelling and a standing wave

Fig. 3 shows the Fourier analysis of the 3D beat across the population. At the comoving frame, Fig. 3(a), the amplitude of static mode in both directions, (*y_c_, z_c_*), are very small, top-row Fig. 3(b,c) due to the beating symmetry. Moreover, the amplitude (middle-row) and phase (bottom-row) of (*y_c_, z_c_*) dynamical modes in Fig. 3(b,c) are analogous, capturing the transversal symmetry, and thus beating isotropy at this frame of reference due to the coordinated sperm rolling (compare SI-Figs. 15,16). Indeed, the travelling wave characteristics of both *y_c_* and *z_c_* coordinates (calculated from phase in Fig. 3(b,c) bottom-row, see Methods) are similar across the population, averaging a frequency of 4Hz, a wavelength of 100*μ*m and a wavespeed of 400*μ*m/s.

**FIG. 3.**
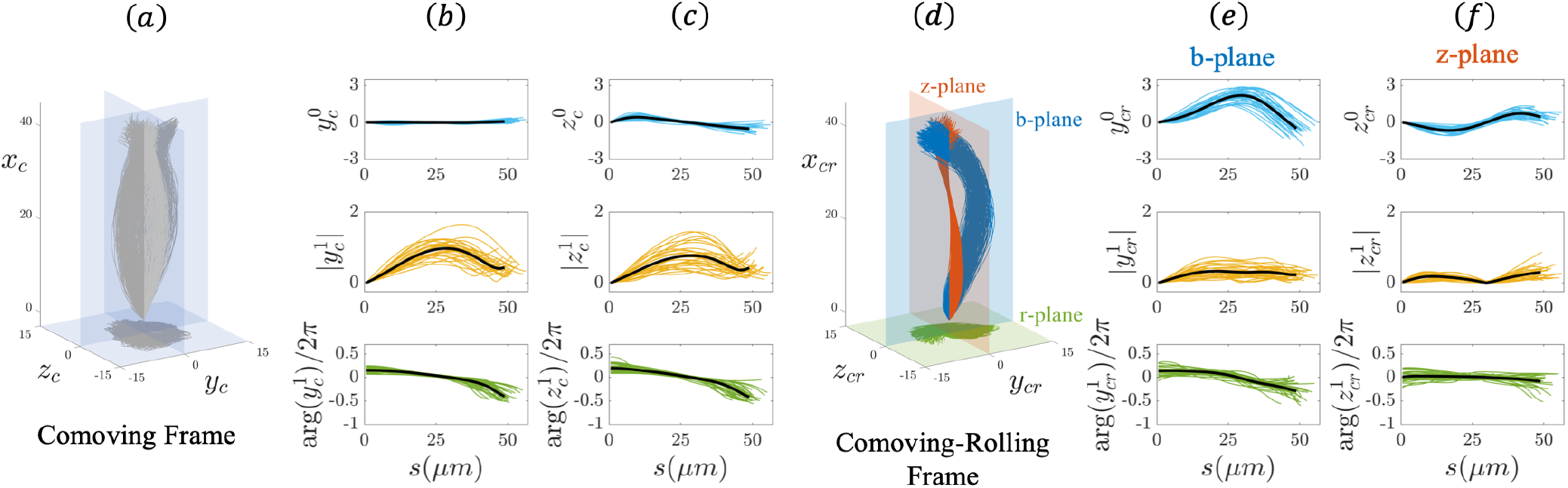
The 3D flagellar beating is a transverse superposition of a travelling and a standing wave. Fourier analysis of each transverse-plane across the sperm population: (a,b)-columns, for the comoving frame coordinates (*y_c_, z_c_*); (c,d)-columns, for the comoving-rolling frame coordinates, (*y_cr_, z_cr_*), denoted, respectively, by b-plane (c) and z-plane (d). Top and middle-rows show the amplitude of the static and dynamic modes, respectively, as a function of arclength. Bottom-row show the phase of the dynamical mode as a function of arclength. Black curves depict averages in the sperm population. The transformations between referential frames and Fourier analysis are detailed in Methods.

At the comoving-rolling frame, Fig. 3(d), the large, but distinct, static modes of b-plane and z-plane show that the beating has a conserved anisotropy across the population, top-row in Fig. 3(e,f). The static mode of the b-plane, *y_cr_*, is highly asymmetric, skewed to positive values, top-row in Fig. 3(e), while the z-plane, *z_cr_* oscillates symmetrically along the arclength in a sinusoidal fashion, top-row in Fig. 3(f). Both (*y_cr_, z_cr_*) static modes are well characterised by a sum of two sines by curve fitting with the population mean (black curves, top-row in Figs. 3(e,f)), respectively, given by 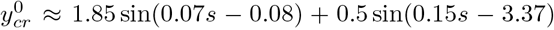, and 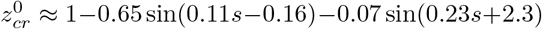 where s is arclength. The amplitude of dynamical mode 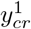 at the b-plane, increases until plateauing, middle-row in Fig. 3(e), while the dynamical mode 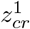 at the z-plane is non-monotonic along arclength, middle-row in Fig. 3(f). The latter is due to the fact that the mid-flagellum is the reference point capturing the flagellar rolling in the comoving-rolling frame of reference (Methods), therefore, by definition, *z_cr_* = 0 at this point in arclength.

The phase of the dynamical mode of *y_cr_*, 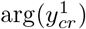, decreases along arclength, bottom-row in Fig. 3(e). Phase differences along flagellar length are expected for propagating undulations, whilst the rate of decay of the phase captures the wavespeed (Methods). Thus the wavespeed of ycr is weakly nonlinear along arclength (see also SI-Fig. 17). The travelling wave characteristics of *y_cr_* (b-plane) average, across the population, a frequency of *ω_y_cr__* = 8Hz, wavelength of 145*μ*m and wavespeed of 1120*μ*m/s. On the other hand, *z_cr_* (z-plane) oscillates with an average frequency of *ω_z_cr__* = 6Hz, wavelength of 1526*μ*m and wavespeed of 5174*μ*m/s, across the population. Such high values of wavespeed/wavelength are due to the almost constant arclength-profile of 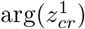, bottom row in Fig. 3(f) and SI-Fig. 12. Small changes in phase across long flagellar distances require very rapid propagation speeds, thus the z-plane wave behaves, effectively, as a standing wave pulsating in time, as also shown in SI-Fig. 18. The flagellar waveform in 3D is thus a superposition of two anisotropic transversal waves, not too dissimilar from electromagnetic waves travelling in space. Here, however, the each transversal wave (*y_cr_, z_cr_*) is a sum of a static and a dynamical mode: an asymmetric travelling wave along the b-plane (blue), and a symmetric standing wave in the z-plane (red), Fig. 3(d) and SI-Figs. 17,18.

### The solid boundary modulates the beat anisotropically in 3D

The presence of the nearby surface decreases the amplitude of the wave propagation in 3D, Fig. 4 and SI-Figs. 11, 12. At the comoving frame of reference, Fig. 4(a,b), the amplitude of *both* the (*y_c_, z_c_*)-dynamical modes are reduced towards the end piece for sperm near to coverslip, whilst the static modes remained unchanged. The dynamical mode of zc is only slightly smaller than yc in Fig. 4(a,b). This is in contrast to the symmetric profiles of both (*y_c_, z_c_*)-dynamical modes for spermatozoa found away from the boundary (red curves, middle-plots in Fig. 4(a,b)). At the comoving-rolling frame of reference in Fig. 4(c,d), however, the influence of coverslip is distinctively anisotropic, as it only affects *one* beating direction, the b-plane - this is seen for both the static and dynamic modes of ycr (Fig. 4(c)). In contrast, the z-plane *z_cr_* remains unchanged for cells swimming near to the coverslip (Fig. 4(d)). In all cases, the phase is weakly perturbed by the solid boundary, bottom-row in Figs. 4(a-d).

**FIG. 4.**
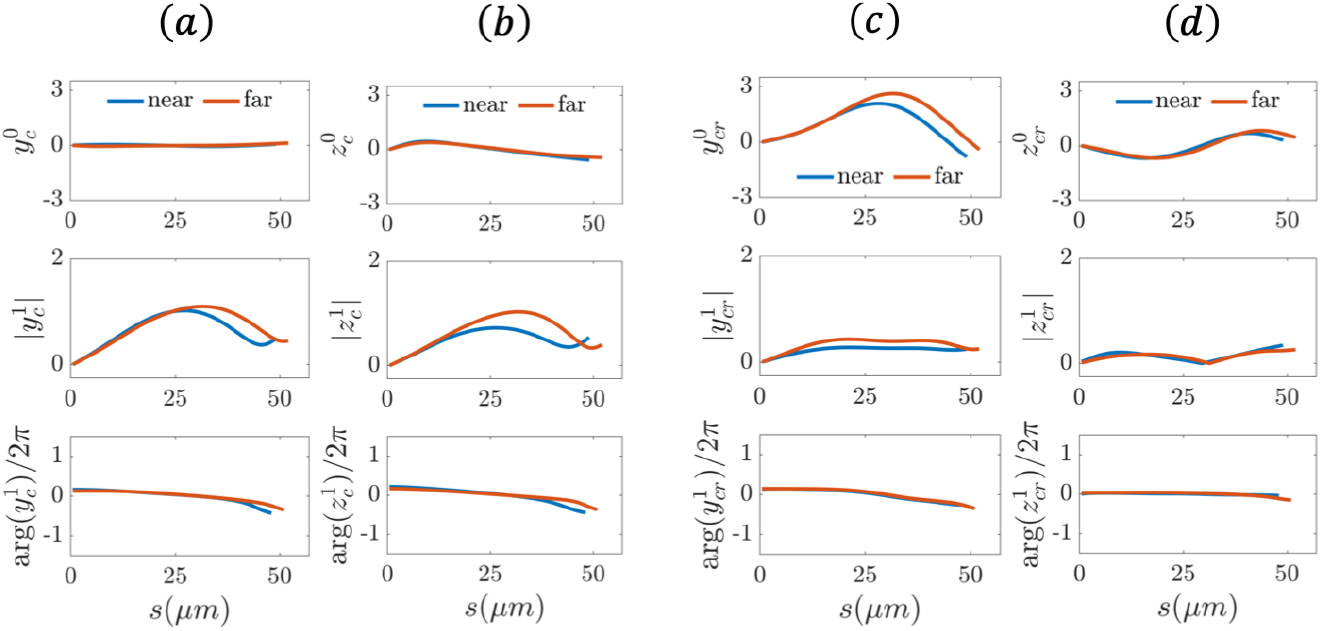
The coverslip modulates the beat anisotropically in 3D. Each column (a-d), top to bottom, respectively, the averages across the population of the static and dynamic modes, and phase of *y_c_* (a), *z_c_* (b), *y_cr_* (c) and *z_cr_* (d) for the sperm near to (blue) and far from (red) the coverslip.

### The static mode reminisces a logarithmic helix and governs the sperm rolling bi-directionality

The shape of the static mode at the comoving-rolling frame is conserved across the population, and define an off-centred, right-handed helix in 3D, **h**(*s*), black curve in Fig. 5(a). The projection of the helix at the rolling plane (green plane) is a counterclockwise-spiral which lacks polar symmetry, i.e. it is skewed to one side (grey projection in Fig. 5(a)). In other words, the helix “unfolds” the spiral along the rolling axis (intersection between the blue and red planes in Fig. 5(a)). The radius of the spiral *r* is a non-monotonic function of both arclength *s* and the polar angle *α* in Fig. 5(b,c). The slope of *r*^2^, before and after *r* = *r_max_* (Fig. 5), is approximately constant and symmetric, as shown by the linear square-markers in Fig. 5(c). The spiral around *r_max_* is thus well approximated by a superposition of two Fermat spirals, with positive and negative slopes, respectively, before and after the maximum value, with *r*^2^ ≈ ±*a*^2^*α* and *α*^2^ = 16.89. The spiral radius, however, has a Gaussian distribution around *r_max_* reflecting its non-monotonicity, with *r*(*s*) = 2.2 exp [−((*s* − 28.12)/16.68)^2^], depicted by the red curve in Fig. 5(b). The spiral static mode is thus reminiscent of logarithmic spirals which are frequently found in nature, with *r* = *ae^bα^* and generic constants *α, b* = 0. However, here, the spiral radius varies non-monotonically, increasing/decreasing at a higher rate than logarithmic spirals in nature.

**FIG. 5.**
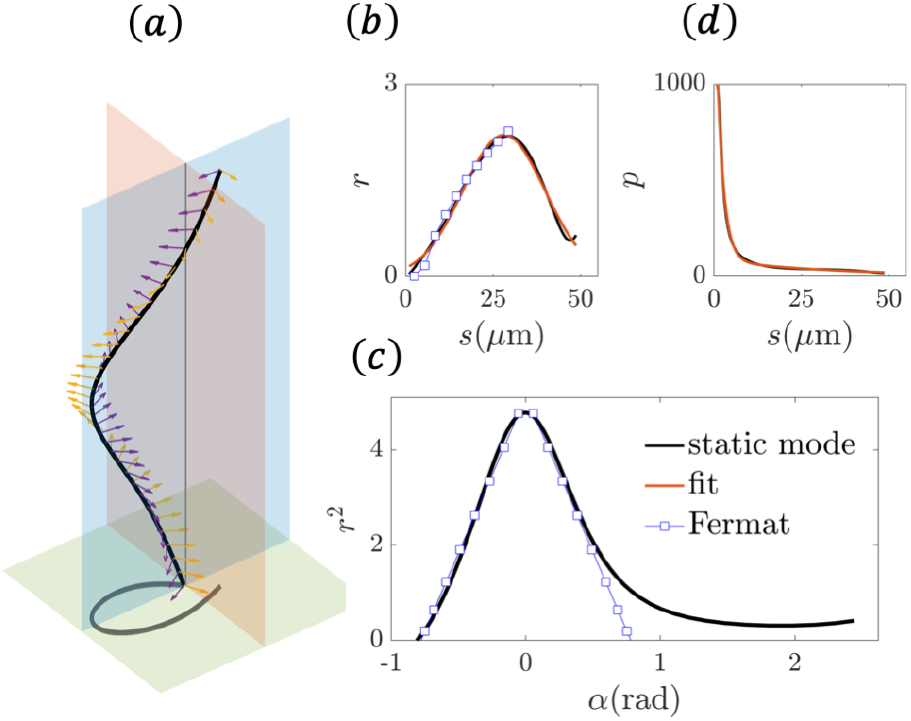
The static mode is a logarithmic-like helix that dictates the sperm rolling direction: (a) The averaged static mode of the coordinates 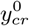 and 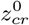 across the population, shown in Figs. 3(c,d) top-row by the black curves, define a right-handed helix in 3D (black curve), denoted by **h**(*s*). The helix projection at the rolling plane (green plane) is an asymmetric, counterclockwise spiral (grey curve). The b-plane and z-plane are shown, respectively, by the blue and red planes. Arrows depicting the Frenet basis (normal and binormal vectors) showing that the helix is right-handed (Methods). The dashed straight-line is the rolling axis; (b) radius of the spiral *r* in polar coordinates (*r, α*) as a function of arclength s (black curve); (c) radius-squared of the spiral, *r*^2^, in function the polar angle *α* (black curve), with blue squares showing the linear regression before and after the maximum at *α* = 0, denoted by *r_max_*. The red curve and blue squares in (b) show, respectively, the Gaussian fit and the linear regression, as in (c), but only for the first-half before *r_max_*; (d) pitch of the helix, *p*(*s*), as a function of arclength *s* (black curve) and its exponential fit (red curve), see main text for details.

The right-handed, logarithmic-like helix in Fig. 5(a) can be expressed in terms of its radius *r*(*s*) and pitch *p*(*s*), as *h*(*s*) = (*h_x_, h_y_, h_z_*) ≈ (*s, r* cos(*sl*), *r* sin(*sl*)) via the wave-number *l*(*s*) = (*r*^2^ + *p*^2^)^-1/2^. The pitch *p*(*s*), Fig. 5(d), decays exponentially along the flagellar length, according to *p*(*s*) = 1651 exp(−0.53*s*) + 84exp(−0.03*s*). Any sign change in either *h_y_* or *h_z_* coordinates causes the handiness of the spiral to switch to a clockwise-spiral, generating instead a left-handed helix. All freely swimming spermatozoa generated counterclockwise-spirals, and thus right-handed helices for their static modes, as in Fig. 5(a). Only two spermatozoa were observed with clockwise-spirals, though with shapes similar to Fig. 5(a) under sign change *h_z_* → −*h_z_*. Incidentally, these two cells (spermatozoa sp29 and sp30 in SI text) were prevented to swim forward by obstacles in their path, though still able to rotate around the rolling axis (see SI video), as in the experiments performed by Ishijima et al. [56]. In all cases, the handiness of the spiral correlated with the sperm rolling direction: counterclockwise-spiral for clockwise rolling (when seen from the posterior end), and clockwise-spirals for counterclockwise rolling, also further discussed in SI text.

### The 3D flagellar curvature and the waveform perversion phenomenon

The 3D curvature, *κ*, and torsion, *τ*, of the flagellum centreline was measured spatiotemporally from experiments (Methods). Fig. 6(a,b) shows a complex train of travelling waves as the flagellum rotates around its rolling axis. Bending waves travel linearly along the flagellum (black slope in Fig. 6(d)) with a non-monotonic amplitude along the arclength, characterised by an abrupt increase at the midpiece and the distal regions, see static mode of *κ* in Fig. 6(h). The waveform torsion is characterised by sharp travelling turns along arclength, Fig. 6(b,c), with co-existing positive and negative turns at the same instant, see blue and yellow ribbons in Fig. 6(c), or equally, the propagating yellow and blue peaks in Fig. 6(e). The helical shape of the flagellum *centreline* undergoes a ‘perversion’ phenomenon by which sections of opposite chirality (handiness) along the flagellum coexist [34, 50, 57]. Here, however, the flagellar sections of opposite chirality travel during the beat, Fig. 6(c,e). The travelling waves of torsion, or rather perversion, propagate with the same speed as the wave of curvature (see linear decay in Figs. 6(h,i) bottom-rows), however, with the manifestation of phase jumps in torsion (sudden changes from blue to yellow in Figs. 6(e)) and time-delays between the wave propagation of *κ* and *τ*.

**FIG. 6.**
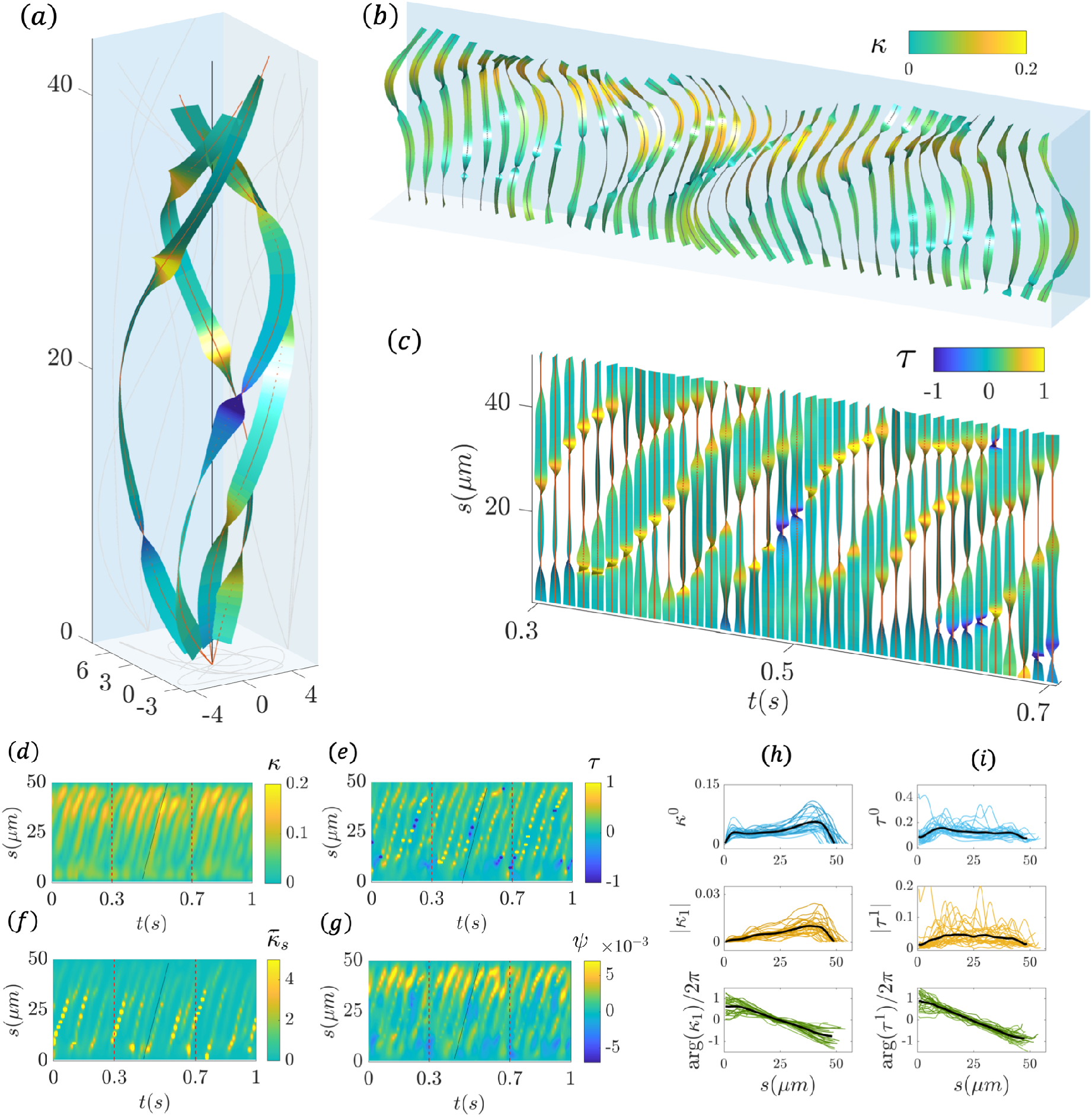
Helical waves of flagellar perversion. (a) typical flagellar bending around the rolling axis (black line) with ribbon’s colour showing the torsion intensity (Methods). (b) the 3D waveform time-series with ribbon’s colour showing the magnitude in curvature. (c) Torsional waves in absence of bending deformation with ribbon colours indicating the torsion. (a-c) all ribbons indicate the torsional angle of rotation (Methods). (d-g) kymographs of the curvature *κ*, torsion *τ*, spiral’s curvature *κ_s_* (projection in the rolling plane) and the chirality *ψ* as a function of arclength *s* and time *τ*. Overlaid black line indicate the speed of propagation is the same in (a-d). The dashed line in (d-g) show the time-duration depicted in (b,c). (h,i) top to bottom: amplitude of the static and dynamic modes, and phase, respectively, for (h)-column the curvature and (i)-column the absolute torsion |*τ*|, with black curves depicting averages in the sperm population. The waveform parameters and Fourier analysis are detailed in Methods.

The curvature of the spiral *κ_s_* in Fig. 6(f), defined as the waveform projection at the rolling plane (Methods), is a linear propagating wave of peaks, similar to the torsional waves (compare the straight-lines in Fig. 6(e,f)). This is due to the formation of sharp kinks in this projection. Incidentally, *κ_s_* propagates similarly to *κ* (compare Figs. 6(d,f)). Large torsion occurs when either *κ*, or rather *κ_s_*, approaches zero, as the centreline torsion scales with 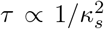. In other words, when curvature is small, the torsion is large, if the out-of-plane component of the waveform is non-zero, leading to the observed phase-lag between curvature and torsion in Fig. 6(d,e). The flagellar perversion is propagated with wave characteristics similar to the curvature (compare Figs. 6(g) and (d)). Nevertheless, the local chirality is not conserved, as indicated by the change in handiness (sign changes) in SI-Fig. 22.

The static and dynamic modes of *κ* and |*τ*| across the population are shown in Fig. 6(h,i). The curvature static mode (top-row in Fig. 6(h)) increases non-monotonically and has two local maxima. The absolute torsion static mode (top-row in Fig. 6(i)) also has a rapid increase at midpiece from which the amplitude decays. The curvature dynamic mode (middle-row in Fig. 6(h)) increases linearly before reaching a maximum at distal end, while the absolute torsion dynamic mode (middle-row in Fig. 6(i)) is parabolically distributed along the length. Curvature and torsion average similar wave characteristics, with a frequency of 14Hz (16Hz), wavelength of 38*μ*m (31*μ*m) and wavespeed of 415*μ*m/s (452*μ*m/s) for the curvature (torsion), as shown in Figs. 6(h,i) bottom-rows. This is expected from the mathematical relation between curvature and torsion (Methods), hence the strong correlation between the beating frequency, defined by the 1st peak of the Fourier spectrum of the curvature *ω_κ_* (arrow in Fig. 7(d)), and the torsion’s frequency *ω_τ_*, as shown in Fig. 7(a) with R= 0.69. Most interestingly, the torsion’s frequency *ω_τ_* displayed the strongest correlation with the sperm rolling frequency, *ω_roll_*, R= 0.71 in Fig. 7(b).

**FIG. 7.**
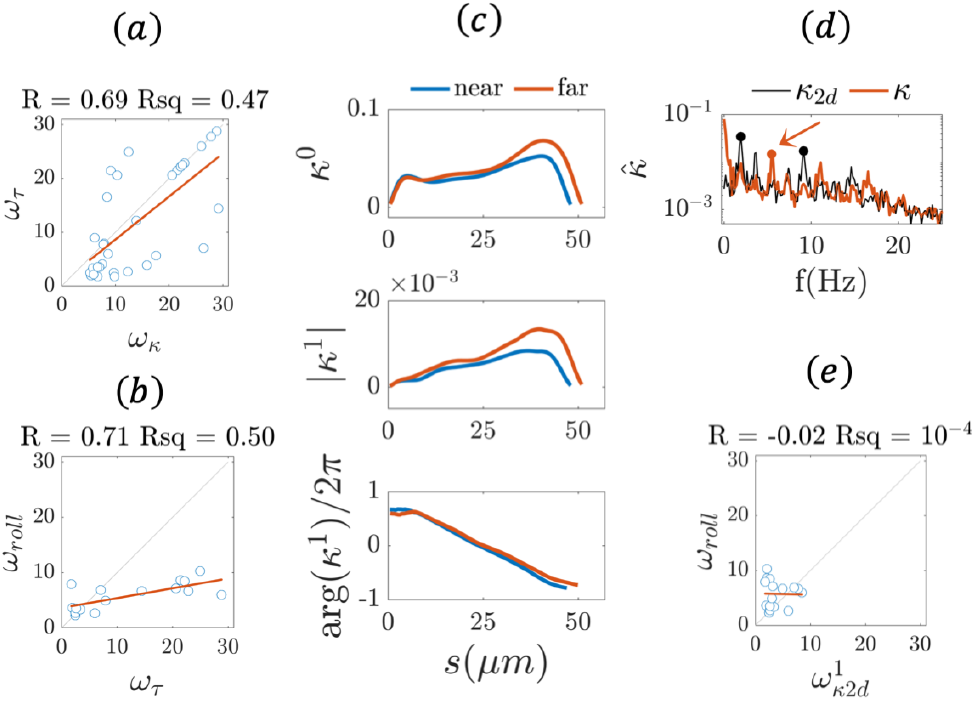
Beating frequency correlations and hydrodynamic boundary modulation of the curvature. (a) correlation between the beating frequency, defined by the 1st peak of the Fourier spectrum of the curvature (arrow in (d)), *ω_κ_*, and the torsion’s frequency, *ω_τ_*. (b) Strong correlation between torsion *ω_τ_* and the sperm rolling frequency *ω_roll_*. (c) top to bottom, respectively, the averages across the population of the static and dynamic modes, and phase of the curvature *κ*, for the sperm near to (blue) and far from (red) the coverslip. (d) Typical Fourier spectrum of the 3D curvature *α*(red) and the 2D curvature *K*_2*d*_ (black), obtained by projecting the 3D waveform in the *xy*-plane (grey curves in Fig. 1(c,d) and Methods). The spectrum of *K*_2*d*_ is characterised instead by two frequency-peaks (black markers), denoted by 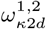. (e) No correlation exist between the 1st frequency-peak of 2D curvature, 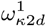 and the sperm rolling *ω_roll_*.

The proximity to the coverslip modulates the beat in 3D by reducing the amplitude of both static and dynamic modes of the curvature (Fig. 7(c)) towards the distal end across the population (blue curves at top and middle-rows in Fig. 7(a) and SI-Fig. 13), similarly to *y_cr_* in Fig. 4(c). The phase of *κ* is equally affected, as show in Fig. 7(c) at the bottom-row: spermatozoa swimming near to (far from) the coverslip were observed to beat slightly faster (slower), with an average frequency of 15Hz (11Hz), wavelength of 35*μ*m (42*μ*m) and wavespeed of 429*μ*m/s (392*μ*m/s).

### 2D microscope projection fails to capture beating asymmetry

The static mode of the 2D curvature *κ*_2*d*_ is negligible in SI-Fig. 14 and Fig. 7(d), thus no intrinsic waveform bias can be detected with 2D microscope projections. The frequency spectrum is characterised instead by two frequency-peaks (black markers in Fig. 7(d)) rather than a single main frequency-peak observed for the 3D curvature *κ* (arrow in Fig. 7(d) and SI-Figs. 23, 24). The 1st frequency-peak of 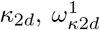, often postulated to be correlated with the sperm rolling frequency, *ω_roll_*, shows no correlation in Fig. 7(e). The 2nd frequency-peak of the 2D curtvature, 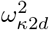, correlates strongly with the beating frequency *ω_κ_* (R= 0.82 in SI-Fig. 1), though *κ*_2*d*_ underestimates wave parameters, and displays a non-existent linear growth of curvature along the arclength (due to the absence of the static mode, SI-Fig. 14). The latter have been the basis of numerous model assumptions and postulations in the literature [10, 24, 26, 36, 58], as discussed in SI text.

## DISCUSSION

We employed high-precision 3D video-microscopy to resolve the rapid movement of the human flagellum in three-dimensions. This allowed, for the first time, the description of the flagellar waveform relative to both the comoving and rolling frame of reference of the sperm flagellum, unveiling the true nature of the waveform unbiased by the sperm rolling. The human sperm flagellum beating, postulated to be symmetrical, is in fact skewed, with a broken left-right symmetry in one plane, with a small, but symmetric out-of-plane motion. The rolling motion thus critically enables spermatozoa to swim in a straight trajectory despite its biased waveform. The latter is achieved via a coordinated 360° degrees rotation of asymmetric bends on one side of the flagellum around the rolling axis, driving symmetric excursions of the flagellum in both *xy* and *xz* directions, Fig. 1(c,d). The rolling-of-asymmetric-bend mechanism induces a sperm kinematic illusion in which the flagellar beating appears to be symmetric for cells swimming in a straight trajectory if observed with traditional 2D microscopy [24, 25, 27, 28, 58], rendering this intrinsic flagellar asymmetry invisible thus far. Similar rolling-of-asymmetric-bend mechanism have also been reported for hamster spermatozoa in which the sperm flagellum induces a rapid side-to-side 180° degrees switch, rather than the gradual 360° rolling for human sperm, to achieve straight-line symmetric swimming despite the asymmetric bending [3].

The movement of the flagellum in 3D is a combination of a planar and out-of-plane motion captured, respectively, by the b- and z-plane components (Fig. 3(d)). The flagellum motion in each beating plane is a superposition of two independent shape modes: a *static mode* whose shape does not change in time, and a *dynamic mode* that oscillates in time (Methods). The waveform of each beating plane share antagonistic properties: the static mode of the b-plane is asymmetric along the flagellum while the z-plane is symmetric (Fig. 3(e,f)). Likewise, the dynamic mode of the b-plane is a travelling wave while the z-plane pulsates as a standing wave (Fig. 3(e,f)). The 3D flagellar beating is thus anisotropic and given by a superposition of two transversal waves: an asymmetric travelling wave and a standing wave. The out-of-plane standing wave in the z-plane further modulates the oscillations between purely planar and weakly non-planar beating in 3D (Fig. 1(g)), therefore critical for the sperm rolling self-organisation in Fig. 1(c,d) and SI-Fig. 3, as no sperm rolling can occur without the out-of-plane component of the beating.

The shape of the static modes define a skewed, righthanded helix whose projection in the rolling-plane is a counterclockwise-spiral reminiscent of logarithmic spirals found in nature (Fig. 5). Generally, the helix is ambidextrous [59], defining a clockwise or counterclockwise spiral (Methods). The spiral shape of the static mode works as a ‘wave guide’ for the propagation of the dynamic mode (Fig. 3(e,f)). As a result, the rolling direction is dictated by the spiral’s handiness, given that travelling waves always propagate from the proximal to the distal direction. The helical flagellum thus rotates around the rolling axis in the opposite direction of the spiral’s travelling wave due to the total balance of momentum in the fluid. Indeed, human spermatozoa with a counterclockwise-spiral (right-handed helix) rolled clockwise (when seen from the posterior end), while clockwise-spirals (left-handed helix) rolled counterclockwise (SI text and SI-Fig. 3).

The waveform torsion is characterised by rapid turns in both directions with phase-jumps and time-delays between distant parts of the flagellum (Fig. 6). This unveiled a new waveform perversion phenomenon manifested via kinks connecting helices of opposite chirality along the flagellum [57]. The travelling waves of perversion induces the lack of local persistent chirality, as previously observed for malaria flagella [50] and human spermatozoa [34], and shown in Fig. 6(g) and SI-Fig. 22 with a finer resolution. Paradoxically, the flagellar trajectories in space define helices with conserved handiness [3, 34, 41, 60], as depicted by the red curve in Fig. 1(a). Indeed, the handiness of the spiral static mode is dominant and dictates the sperm rolling direction (Figs. 3(e,f) and 5). Furthremore, torsional waves are strongly correlated with sperm rolling (R= 0.71, Fig. 7(b)). The latter thus reconciles the current paradigm disconnecting the flagellar chirality from the sperm rolling motion, motivated by earlier observations of the lack of persistent chirality during the beat [34, 50].

Human spermatozoa may achieve a high levels of control of the flagellar helical handiness, and subsequent rolling motion, without requiring complex dynamical regulatory mechanisms. This could be achieved, for example, by simply shaping the static mode with a stable asymmetric binding of molecular motors on one side of the flagellum [1]. Indeed, sign changes of any coordinate-component of the spiral can switch the handiness of the flagellar helix. Likewise, flagellar buckling may also modify the spiral’s handiness and the sperm rolling direction, for example, when spermatozoa are prevented to swim freely due to obstacles, though still free to rotate around its rolling axis [56]. This potentially conciliates contradictory observations thus far showing unidirectional, bidirectional and even intermittent directionality of rolling in human spermatozoa [3, 34, 41, 56, 60] (SI text).

The logarithmic helix of the static mode entails that the forces along the flagellum are non-uniform, asymmetric and helically distributed (Fig. 5), in contrast with the circular arcs in *Chlamydomonas* mutants [8]. Our observations are consistent with recent reports showing a hierarchy of distinct asymmetries in flagellar systems, from asymmetric distribution of molecular motors [1] to observations of asymmetric bending in hamster spermatozoa [3] and asymmetric static modes of *Chlamydomonas* [5, 6], as well as asymmetrical distribution and activity of dyneins [13–15], including an ever growing number of asymmetric regulatory complexes [6, 8, 16–19], in addition to other unknown biases from less well-studied motor proteins [20]. While the exact molecular mechanism underlying the observed asymmetric static shape is still unknown [5, 6], our results show that, together with murine spermatozoa [3] and *Chlamydomonas* flagellum [5], the human sperm flagellum is part of the group of flagellates that exhibit an intrinsically asymmetric static mode, potentially reflecting an universal asymmetry of the axoneme conserved across these species.

The wave amplitude in the b-plane is much larger than in the z-plane, which posses instead a sinusoidal distribution in arclength (Fig. 3(e,f), top-row). In other words, the waveform is anisotropic, as depicted in Fig. 8(a). This may find its origin on the ultrastructural complex reinforcing the axoneme in human sperm [10, 12], represented in Fig. 8(b). Outer dense fibres are connected to the microtubule doublets forming the 9 + 9 + 2 structure. The outer dense fibres 8 and 3, however, are replaced by two diametrically opposite longitudinal columns (LCs) at the principal piece, as shown in Fig. 8(c). Thus, this introduces a mechanical anisotropy along the flagellum: it suppresses bending in one direction and facilitates in the other. The large bending observed in one direction could be facilitated by the ‘weaker’ side of the flagellum, so that the b-plane would lie perpendicular to the LCs (blue double arrow in Fig. 8(c)). On the other hand, bending could be hindered by the ‘stronger’ side of the flagellum, so that the z-plane would lie parallel to LCs, as illustrated in Fig. 8(c). Coincidently, upon increased hydrodynamic friction (for sperm swimming near to the coverslip), only modest changes in wave amplitude were observed at the z-plane (stronger side of the flagellum), and Fig. 4(d), while the b-plane (weaker side) was greatly affected by the nearby surface in Fig. 4(c). Finally, the sinusoidal shape of the static mode in the z-plane (Fig. 3(f), top-row) could also be a manifestation of another structural effect, for example, via the axonemal counterbend phenomenon, in which proximal bending causes the distal end to bend in the opposite direction, resembling the sine-like shape in Fig. 3(f) top-row [11, 61, 62].

**FIG. 8.**
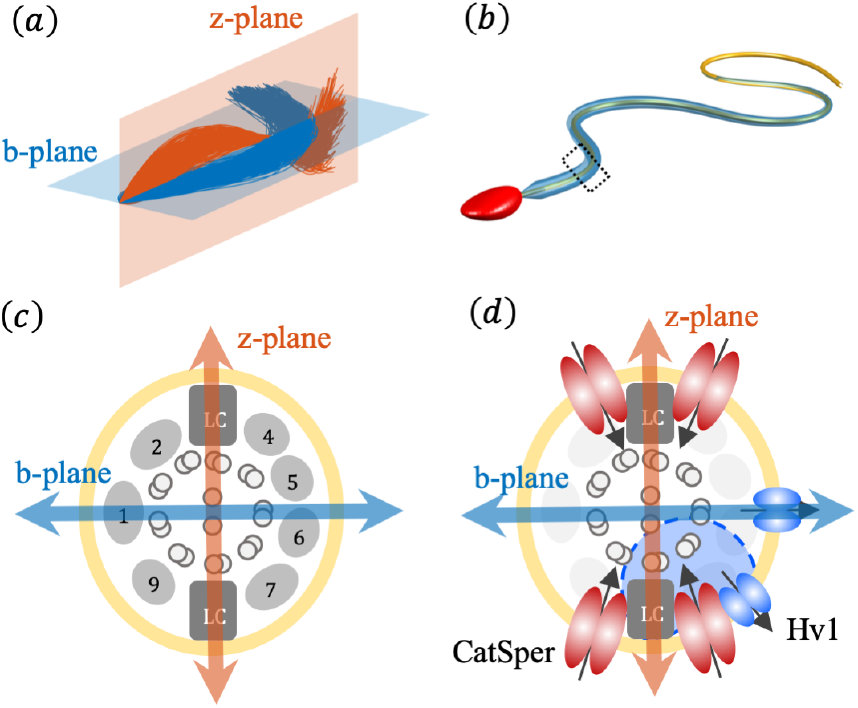
Ultrastructural and molecular origin of the flagellar anisotropy: (a) The flagellar beating anisotropy in 3D, as in Fig. 3(d), showing the b-plane and z-plane. (b) Representation of a mammalian sperm flagellum showing the tapering of the reinforcing ultrastructure along the flagellar length (blue), with the axoneme depicted in yellow. (c) depicts the crosssection of a mammalian sperm flagellum (rectangle in (b)) showing that the 9 + 9 + 2 ultrastructure [10] is anisotropic. The 9 + 2 axoneme is surrounded by 9 outer dense fibres (gray), however, the outer dense fibres 8 and 3 are replaced by, two diagonally opposite, longitudinal columns (LC). The LCs constrain the beating to “weaker” side of the flagellum (blue doable-arrow), thus potentially co-aligned with the b-plane in (a). The reduced amplitudes of the z-plane (red) in (a) are likely to be suppressed by the “stronger” side of the flagellum, and thus aligned with the plane defined by the LCs (red doable-arrow). (d) Anisotropic bilateral localisation of Hv1 proton channels (blue eclipses) compared with the symmetrical, quadrilateral localisation of CatSper ion channels (red ellipses) reported by Lishko’s group [17], in a flagellar cross-section. Dashed circular region (light blue) shows the CatSper channel mostly affected by one of the Hv1 channels in close proximity, which is also located near to one of the LCs, thus affecting molecular motors anisotropically along the “stronger” side of the flagellum.

Each beating plane is dynamically very distinct: the wavespeed of the z-plane is five times higher, and wavelength is tenfold larger, than the travelling waves of the b-plane. Thus, the z-plane effectively pulsates as standing wave (bottom-row in Fig. 3(e,f) and SI-Fig. 18). This could be indicative of an independent, though anisotropic, regulatory mechanism modulating each plane; or else, this would require molecular motors alone to work differently in each beating plane. Furthermore, the standing waves in the z-plane are unlikely to be achieved via molecular motor dynamics alone as this would require a fast, synchronised motor coordination over long flagellar distances, not yet reported in the literature. Indeed, molecular motor self-organisation is manifested through time-delayed oscillations along the flagellum [38, 63, 64] which leads to the generation of propagating waves, and therefore the most likely mechanism to govern the asymmetric travelling waves observed in the b-plane (Fig. 8(a)) [1, 2, 5].

The standing wave pulsation of the z-plane indicate the existance of a fast regulator along the flagellum length (Fig. 3(f)). Ion channels, such as CatSper and Hv1, are docked at the plasma membrane to ensure fast signal transduction in the whole flagellum (Fig. 8(d)), and are critical for hyperactivation, rheotaxis and sperm rolling [17–19, 34]. Furthermore, species specific adaptation could also lead to specialised localisation of these regulatory complexes. For example, the human sperm flagellum has the largest Hv1 current density proton channel of all cell types [17, 18]. However, unlike CatSper channels [19] which possess a symmetrical quadrilateral localisation at the flagellar cross-section, Hv1 is distributed asymmetrically and unilaterally along the flagellum [17], and may be related with the observed pulsation in the z-plane, as indicated in Fig. 8(d).

The z-plane modulates the out-of-plane component of the beat (Fig. 8(a)), its quasi-planarity and subsequent sperm rolling self-organisation. Indeed, upon inhibition of Hv1, Miller et al. [17] demonstrated that the 360° rolling of human sperm was dramatically decreased, often leading to a partial 180° rolling events, much similar to the motion of murine sperm [3, 19], which indeed lack Hv1 expression [18]. As such, given the observed correlation between the sperm rolling and the waveform torsion in Fig. 7(b), the Hv1 channel may play an important role while regulating the waveform torsion. Furthermore, Miller et al. [17, 18] showed that Hv1 may alkalinize a sub-portion of the axoneme and up-regulate a sub-set of CatSper channels in close proximity (blue region in Fig. 8(d)). This would result on a local increase in Ca^2+^, causing the arrest of dyneins and increasing the effective rigidity longitudinally within the flagellum. Most strikingly, the Hv1 channel likely up-regulates the CatSper located near one of the LCs (dashed circle in Fig. 8(d)), thus potentially only affecting one beating plane, the z-plane (red double arrow in Fig. 8(d)). This indicates that, once again, the z-plane could be co-aligned with the plane composed by the LCs, similarly to the mammalian ultrastructure localisation in Fig. 8(c). Thus the correlation between the localisation of the LCs and Hv1 channels, and the orientation of the z-plane in Fig. 8(d) might not be incidental.

Even though it is not possible to exactly assert whether the Hv1 channels are responsible or not for the pulsations of the z-plane observed here, our results show that symmetry can be recovered by anisotropically controlling asymmetric bends within an intrinsically asymmetric system. The rolling-of-asymmetric-bend control cancels out the umbrella of intrinsic asymmetries present at the molecular level in the sperm flagellum. This is achieved by exploiting the third-dimension to induce sperm rolling, symmetric excursions of a biased waveform, and subsequent straight-line swimming [24]. While large asymmetric waves contributes to the overall cell propulsion, the sperm rolling self-organisation may also regulate the swimming direction, by exploiting fast or slow rotations of asymmetric bends accordingly. This is in contrast with 2D microscopy which introduces artefacts and does not capture the intrinsic anisotropy and asymmetry of the beat; therefore, it is necessary to reassess observations of symmetric beating in 2D for rolling spermatozoa, and observations of asymmetric waveforms, such as during sperm hyperactivation, as discussed in SI text. The 3D imaging and mathematical analysis employed here may prove an essential tool for future studies on the functional nature of both molecular motor asymmetric dynamics and anisotropic ion channel orchestration leading to flagellar self-organisation.

## METHODS

### Sperm preparations

Human ejaculates were obtained from healthy donors by masturbation after at least 48 hours of sexual abstinence. Written informed consent forms were signed by all donors. All human semen samples fulfilled the requirements determined by the World Health Organisation. Highly motile sperm were recovered after a swim-up separation for 1 hour in Ham’s F-10 medium at 37°C in a humidified atmosphere of 5% CO2 and 95% air. Sperm cells were centrifuged 5 min at 3000 rpm and resuspended in physiological salt solutions at approximately 107 cells/ml. The physiological solution was in mM: 94 NaCl, 4 KCl, 2 CaCl2, 1 MgCl2, 1 Na Piruvate, 25 NaHCO3, 5 Glucose, 30 HEPES, 10 Lactate at pH 7.4.

### 3D imaging microscopy

A piezoelectric device P-725 (Physik Instruments, MA, USA) mounted between a 60X 1.00 NA water immersion objective (Olympus UIS-2 LUMPLFLN 60X W) and an inverted microscope Olympus IX71, also mounted on an optical table (TMC), was used. The piezoelectric device oscillates due to a servo-controller E-501 via a high current amplifier E-505 (Physik Instruments, MA, USA). The servo-controller is triggered to vibrate by a ramp signal from the E-506 function generator. A synchronising TTL pulse from the servo-controller triggers the high speed camera NAC Q1v with 8 Gigabyte RAM (recording up to 3.5 seconds at 640 x 480 pixels at 8,000 images per second). Temperature was controlled with a thermal controller (Warner Instruments TCM/CL-100) with a chamber Chamlide (CM-B18-1) (details in Corkidi et al. [40, 47]).

### Flagella segmentation in 3D

The 3D centreline of the flagellum coordinates were obtained for each 3D stack using a semi-automatic algorithm. For each 3D stack, the flagellum distal point is manually selected from which an automatic algorithm detects the 3D path connecting the sperm head and the selected distal point (Hernandez-Herrera et al. [49]). The centroid of the sperm head is used as the initial point to trace the flagellum (3D voxels coordinates), guided by a cost function inversely proportional to the image intensity using a minimal path algorithm, providing the flagellum centreline in 3D voxel coordinates. The 3D spatial coordinates are obtained directly from the microscope pixel resolution and the position of piezoelectric device. The extracted position of the flagellar coordinates in respect to the laboratory fixed frame of reference, **X**(*s, t*) = (*x, y, z*), are then parametrised by arclength s with a total of 100 discretization segments. Data smoothing routines using the MATLAB ‘csaps’ function were used to remedy random loss of the distal flagellum, and remove spatial and temporal noise as detailed in [22]. The smoothed data were compared post-hoc with the original data to ensure that this process did not cause significant deviations from the original data.

### Human sperm near to and far from the coverslip

In order to investigate the potential for asymmetric, hydrodynamic influence and modulation of non-slip boundaries on the flagellar waveform in 3D, we considered swimming spermatozoa at distinct heights from the bottom coverslip totalling 30 cells with heights between 0 to 85 *μ*m. Spermatozoa heights above 20*μ*m are defined as ‘far from’ the surface (sp20 to sp28 in SI-Figs.), whilst ‘near to’ cells have heights below 20*μ*m (sp1 to sp19 in SI-Figs.). In addition to those, two sperm cells were observed to not swim freely due to obstacles in their paths (sp29 and sp30), and thus they were excluded from the population averages involving free swimming sperm. sp29 and sp30 though prevented to swim freely, they were able to rotate around the rolling axis, see SI-Fig. 3 and SI video.

### Inertia ellipsoids of the flagellum

The total moment of inertia ellipsoid of the flagellum shape is calculated directly from waveform shape **X**(*s, t*), as proposed by [34]. from which eigenvalues and eigenvectors are calculated to provide the orientation and dimensions of the major, *a*, and minor, *b, c*, axis of inertia ellipsoid for each time-frame. The ratio *c/a* of the inertia ellipsoid thus capture the three-dimensionality of the flagellum in space (Fig. 1(g)), or rather the planarity of the waveform, as if *c/a* = 0 the flagellar waveform is a completely planar curve. The orientation of the major axis of the inertia ellipsoid relative to the solid boundary, located at the plane *z* = 0, defines the angle of attack of the flagellum shape at each instant (Fig. 1(h)).

### The comoving and rolling frame of reference

The 3D flagella coordinates relative to the laboratory fixed frame of reference, **X**(*s, t*) = (*x, y, z*), is transformed into the comoving frame of reference **X***_c_* = (*x_c_, y_c_, z_c_*), by first translating the waveform relative to the head position and then rotating the waveform around *z*-axis so that the mean swimming axis is conveniently aligned with the *x*-axis (SI-Fig. 2). The comoving frame **X***_c_* is thus the frame of reference translating with the sperm head and rotating with the main swimming axis, as shown in Figs. 1(c,d) and SI-Fig. 4. 3D excursions of the flagella beat induces sperm rolling around the mean axis, defining in this way the ‘rolling axis’ of the flagellum, denoted by the black curve in Figs. 1(c,d). The comoving frame of reference is transformed into the comoving-rolling frame of reference **X***_cr_* = (*x_cr_, y_cr_, z_cr_*) by simply rotating the waveform around the rolling axis, taking the angular displacement of mid-flagellar section at the rolling plane as the reference rolling angle (red plane in Figs. 1(c,d)). In other words, no sperm rolling occurs in this reference frame. By construction, the mid-flagellar section of the flagellum is always positioned at the *z_cr_* = 0 with *y_cr_* > 0 at the rolling plane (green plane in Fig. 2(e) and Fig. 3(d)). SI videos show a typical flagellar beat as seen from: the laboratory fixed frame, the comoving frame and the comoving-rolling frame of references, respectively. The kymographs of the *y* and *z* coordinates in arclength and time for each spermatozoa are shown in SI-Figs. 15-18 for both frames of reference.

### Principal component analysis of the 3D waveform

We employed principal component analysis (PCA) on the 3D flagellar shape at the comoving, **X***_c_*(*s, t*), and comoving-rolling, **X***_cr_*(*s, t*), frames of reference. For this, we generalised the previous PCA of scalar quantities [26, 38, 65, 66] to three-dimensional vectorial functions, in which each spatial coordinate represents an spatiotemporal map in arclength and time, (*s, t*). With arclength discretised into m values, *s*_1_,…,*s_m_*, and time discretised into *n* values, *t*_1_,…,*t_n_*, the augmented matrix for the flagellum coordinates in any generic frame of reference is simply the vector **X***_iα_* = [*x*(*t_i_*, *s_α_*), *y*(*t_i_*, *s_α_*), *z*(*t_i_*, *s_α_*)], and its augmented temporal average is 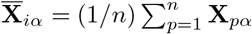. As follows, the eigenvectors of the 3*m* × *m* vectorial co variance matrix, 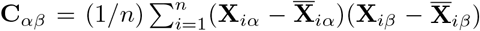, provide a basis for the 3D flagellar coordinates, with m eigenvectors ordered by the size of the associated eigenvalues. Each eigenvector corresponds to a vectorial set of 3D coordinates that define a flagellum shape, also known as PCA modes. Data reconstruction utilising a simple linear combination of the first two PCA modes are depicted in Fig. 2(b,f) for, respectively, the comoving and comoving-rolling frames of reference. Mathematically, the PCA reconstruction of the waveform is simply 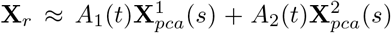, where *A*_1_(*t*), *A*_2_(*t*) are the temporal shape scores for the first 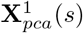 and second 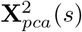 PCA spatial modes, respectively. The contribution of each isolated PCA mode on the course of time, 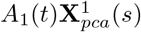 and 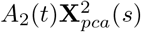, define a surface in space, shown in blue for the 1st PCA mode and in orange for the 2nd PCA mode in Fig. 2(c,g), thus depicting the two principal directions of the flagellum beat in 3D for each frame of reference. The PCA reconstruction, PCA modes and mode amplitudes for each spermatozoa are shown in SI-Figs. 5-10 for both comoving and comoving-rolling frames of reference.

### Fourier analysis and travelling wave parameters

We performed Fourier analysis of the spatiotemporal maps of flagellar coordinates at different frames of references, as well as, for the curvature and torsion of the waveform. In all cases, the input signal is an undulating wave, generically written as *f* = *f* (*s, t*), thus a function of arclength *s* and time *t*. We therefore decompose any generic signal *f* = *f* (*s, t*) as a sum of Fourier modes, 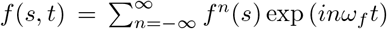, where *ω* is the angular frequency of the signal, with *ω_f_*/2*π* is the signal frequency in cycles per second, *i* is the imaginary number and *f^n^*(*s*) are the spatial modes indexed by *n*. Each mode *f^n^*(*s*) is a complex-valued function of arclength and captures both the amplitude and the phase of the signal along arclength. The 0th mode, *n* = 0 of the Fourier spectrum, is referred as the “static mode” as its time-dependence vanishes, and corresponds in fact to the time-average of the signal *f*^0^(*s*) along arclength - denoted in the main text as superscript 0 of the signal, for example, *κ*^0^(*s*) is the static mode of the cruvature. The highest peak in the Fourier spectrum defines the frequency *ω_f_* of the 1st mode, *n* = 1, also referred as the “dynamical mode”. The spatiotemporal signal is then approximated by a simple linear combination of the static and dynamic modes, and the signal reconstruction reads *f_r_*(*s, t*) ≈ *f*^0^(*s*) + |*f*^1^(*s*)| sin (*ω_f_t* + *ϕ*(*s*)) where |*f*^1^(*s*)| is the amplitude of the dynamical mode and *ϕ*(*s*) = arg(*f*_1_(*s*)) is the phase of the oscillation, from which the wave parameters may be deduced directly (for example, the rate of decay of the phase along arclength captures the wavespeed). Fig. 2(d,h) shows an exemplar of the reconstructed flagellar waveform using the static and dynamic modes of the waveform coordinates, respectively, for the comoving and comoving-rolling frame of references. The black and red curves show, respectively, the individual contributions of the static and dynamic modes (left-plot in Fig. 2(d,h)), and their sum thus provide the reconstructed waveform (middle-plot in Fig. 2(d,h)).

### Waveform curvature, torsion and chirality

The tangent, normal and binormal vectors are calculated directly from flagellum coordinates **X**(*s, t*), respectively, via **T** = **X′**, **N** = **T′**/*κ* and **B** = **T** × **N**, defining the Frenet basis [**T, N, B**] along arclength s at each time *t*, where *κ* = ||**T′**|| and *τ* = −**N** · **B′** are, respectively, the 3D curvature and torsion of the flagellum’s centreline. Primes denote derivatives in arclength, *d/ds*. It is worth noting that the waveform torsion does not capture the mechanical ‘twist’ of the flagellum, but rather the three-dimensional turning of the flagellum centreline in space. Indeed, a purely planar curve embedded in a three-dimensional space has *τ* = 0. Thus, while the curvature of the waveform captures the actual bending of the flagellum in 3D, the waveform torsion measures the kinematic “three-dimensionality” of the centreline of the flagellum. We also define the “waveform spiral” as the shape of the projection of the waveform in the rolling plane (Fig. 3(d)), from which the curvature of the spiral, *κ*_s_, can be derived as above. The chirality or handiness (helicity) of the waveform depends its orientation [59]. For simplicity, the local chiral angle of each section of the flagellum centreline, *ψ*(*s_α_, t*), is defined as the angle between the discrete segment **T***_α_* and the plane defined by **T***_α−2_* × **T***_α−1_* for each flagellar segment *s*_3_,…,*s_m_*, such that if *ψ* is positive, the flagellar section is righthanded, or if negative, the flagellar section is left-handed [50]. The magnitude of the discrete angles *ψ* however depends the number of segments m [67]. The kymographs of the y and z coordinates in arclength and time for each spermatozoa are shown in SI-Figs. 15-18 for both frames of reference. The kymographs of *κ, τ, κ_s_* and *ψ* in arclength and time, for each spermatozoa, are shown in SI-Figs. 19-22.

## ACKNOWLEDGMENTS

GC gratefully acknowledges financial support by Conacyt 253952.

## AUTHOR CONTRIBUTIONS

HG, AD and GC designed research; HG, PHH and FM performed image processing and mathematical data analysis; FM, PHH and GC performed experiment; HG, PHH, FM, AD and GC wrote the paper.

## ADDITIONAL INFORMATION

Supplementary Information accompanies this paper at doi:

